# Managing Autofluorescence in Spectral Flow Cytometry for Macrophage Identification in the Liver

**DOI:** 10.64898/2026.01.28.702253

**Authors:** Sabine Daemen, Panos Barlampas, Xiaodi Zhang, Casper Schalkwijk, Kristiaan Wouters

## Abstract

Autofluorescence (AF) in biological tissues arises from the natural emission of light by intra- and extracellular molecules upon light absorption. Conventional flow cytometry cannot correct for cellular AF, leading to distorted signals and measurement errors. While spectral flow cytometry enables AF visualization and extraction, accurately correcting for AF remains challenging in complex biological samples containing multiple cell types with distinct AF properties, such as the liver. Additionally, pathological processes such as inflammation and fibrosis can alter tissue composition and activate specific cell types, further modifying AF characteristics across experimental conditions. Macrophages are among the most autofluorescent immune cells, exhibiting fluorescence emission across the entire spectrum of light. Recent studies have demonstrated substantial heterogeneity in the phenotypes of resident and recruited macrophages both in the healthy liver and during Metabolic Dysfunction-Associated Steatohepatitis (MASH). Given their critical role in liver disease pathophysiology, we developed a spectral flow cytometry approach to identify and analyze all macrophage subpopulations in healthy and MASH murine livers. Our findings show that healthy, steatotic and MASH livers exhibit distinct and heterogeneous AF signatures. Furthermore, inadequate AF extraction compromised accurate quantification of hepatic macrophages and differentiation of macrophage subsets.

## Introduction

Autofluorescence (AF) in biological tissues is the natural emission of light from intra- and extracellular molecules upon light absorption. Well-known AF substances include NADPH, vitamin A, and extracellular matrix components such as collagen and elastin. Autofluorescence has long hindered the analysis of certain tissue cells, particularly myeloid cells, prompting many efforts to overcome these issues (1, 2). Conventional flow cytometry cannot correct for cellular AF, leading to distorted signals, low staining indices and measurement errors. For instance, FoxP3 expression previously attributed to macrophages was later found to result from their high AF signal (3). Spectral flow cytometry has emerged as an alternative, allowing for the discrimination of an increasing number of immune cell markers. Unlike conventional methods, spectral cytometry enables visualization and subtraction of AF by incorporating the AF signature into unmixing algorithms (4). Nevertheless, adequate AF extraction remains a challenge in biological samples with complex and heterogeneous AF. The AF signature of a sample results from the combined AF signatures of its various cell types. New software tools now facilitate the identification of distinct AF spectra within a sample. In this respect, Kharraz et al. demonstrated that iterative subtraction of multiple AF signals improved the resolution of tissue macrophage populations by incrementally removing individual AF signatures in mouse skeletal muscle (5).

As pathological processes such as inflammation and fibrosis develop, tissue composition becomes increasingly complex, leading to changing and more diverse AF signatures. Meanwhile, advancements in immunology research, such as single-cell RNA sequencing, continue to reveal previously unknown immune cell subsets and activation states with distinct roles in homeostasis and disease. These immune subpopulations can exhibit varying AF signatures depending on their activation state, further complicating their accurate identification in flow cytometry. Importantly, it has been shown for example in the fetal liver, that AF extraction with spectral flow cytometry was able to distinguish cell populations difficult to distinguish with conventional flow cytometry (6).

The liver plays a central role in metabolism and detoxification, leading to the accumulation of numerous AF molecules and making it one of the most autofluorescent tissues. Indeed, hepatic AF emission spans the entire spectrum of light (7). In addition, several pathogenic hepatic conditions, such as steatosis and fibrosis, observed in the development of hepatic diseases including Metabolic dysfunction-Associated Steatotic Liver Disease (MASLD), worsen the AF properties of hepatic cells (8). MASLD is becoming the primary cause of liver disease worldwide as it is strongly associated with obesity. MASLD is characterized by liver steatosis which can further progress to liver inflammation and fibrosis, termed Metabolic dysfunction-Associated SteatoHepatitis (MASH). In mice, the development of steatosis and fibrosis leads to increased AF intensity and shifts in the AF signature of liver tissue (8). Moreover, in humans AF measurements from liver biopsies have been shown to differentiate normal, steatotic and fibrotic livers (9).

An essential immune cell in the pathogenesis of MASLD, and associated liver inflammation and fibrosis, is the macrophage. Recent studies demonstrated substantial heterogeneity in the phenotypes of both resident and recruited macrophages in the healthy liver and during MASH (10), including resident Kupffer Cell (KC) subpopulations and several population of monocyte-derived macrophages, including monocyte-derived KCs (moKCs) and lipid-associated macrophages (LAMs). Spectral flow cytometry has become essential for measuring the growing number of phenotypic markers required to identify and analyze these subpopulations. In particular, identifying LAMs has been challenging with conventional flow cytometry and has mostly relied on negative gating (11). Macrophages are highly phagocytic cells, and previous studies have shown that their phagocytic capacity is linked to their level of AF (12). As such, their AF emission can vary significantly depending on both their activation state and the (patho)biological environment in which they reside.

In summary, two main challenges arise when applying spectral flow cytometry in in highly fluorescent and complex samples like the liver: (1) shifting of AF signatures between experimental conditions and (2) the increasing need to discriminate between small immune cell subpopulations that may present different AF properties, in particular myeloid cells. Here we aimed to set-up a spectral flow cytometry approach to identify and analyze macrophage subpopulations in healthy and MASH murine livers. We compared default AF extraction to a more rigorous approach, where we first identified multiple AF signatures, which were then used as fluorescent tags in unmixing. Moreover, we compared the application of this method in 2 different experimental conditions known to affect AF, i.e. liver steatosis and MASH. We demonstrate that inadequate AF extraction resulted in the inability to quantify hepatic macrophages and to distinguish small macrophage subpopulations. Here, we present a spectral flow cytometry panel and a practical framework for implementing spectral flow cytometry to analyze all recently identified macrophage subsets in healthy and MASH murine livers.

## Results

### Multiple autofluorescence extraction and flow cytometry beads are essential to correctly identify liver myeloid cells

We aimed to develop and optimize a panel of comprehensive myeloid markers for spectral flow cytometry to detect all recently identified macrophage subpopulations in healthy and MASH livers. Initial tests for general optimization of dissociation and measurement procedures included a 7-marker panel (Supplementary Table 1) with commonly used fluorochromes to identify macrophages (F4/80^+^, CD11b^int^), neutrophils (CD11b^+^, Ly6G^+^) and monocytes (CD11b^+^, Ly6C^+^) in the liver. We first tested the use of single stains of the liver cells as reference controls and applied the default AF extraction option available in the SpectroFlo® software during the unmixing. Although a separated macrophage population (CD11b^int^ F4/80^hi^) could be identified, defined populations of Ly6G^+^ neutrophils or Ly6C^+^ monocytes were not detected within the myeloid CD11b^+^ cells (Figure 1A), suggesting unmixing errors.

**Figure 1.**
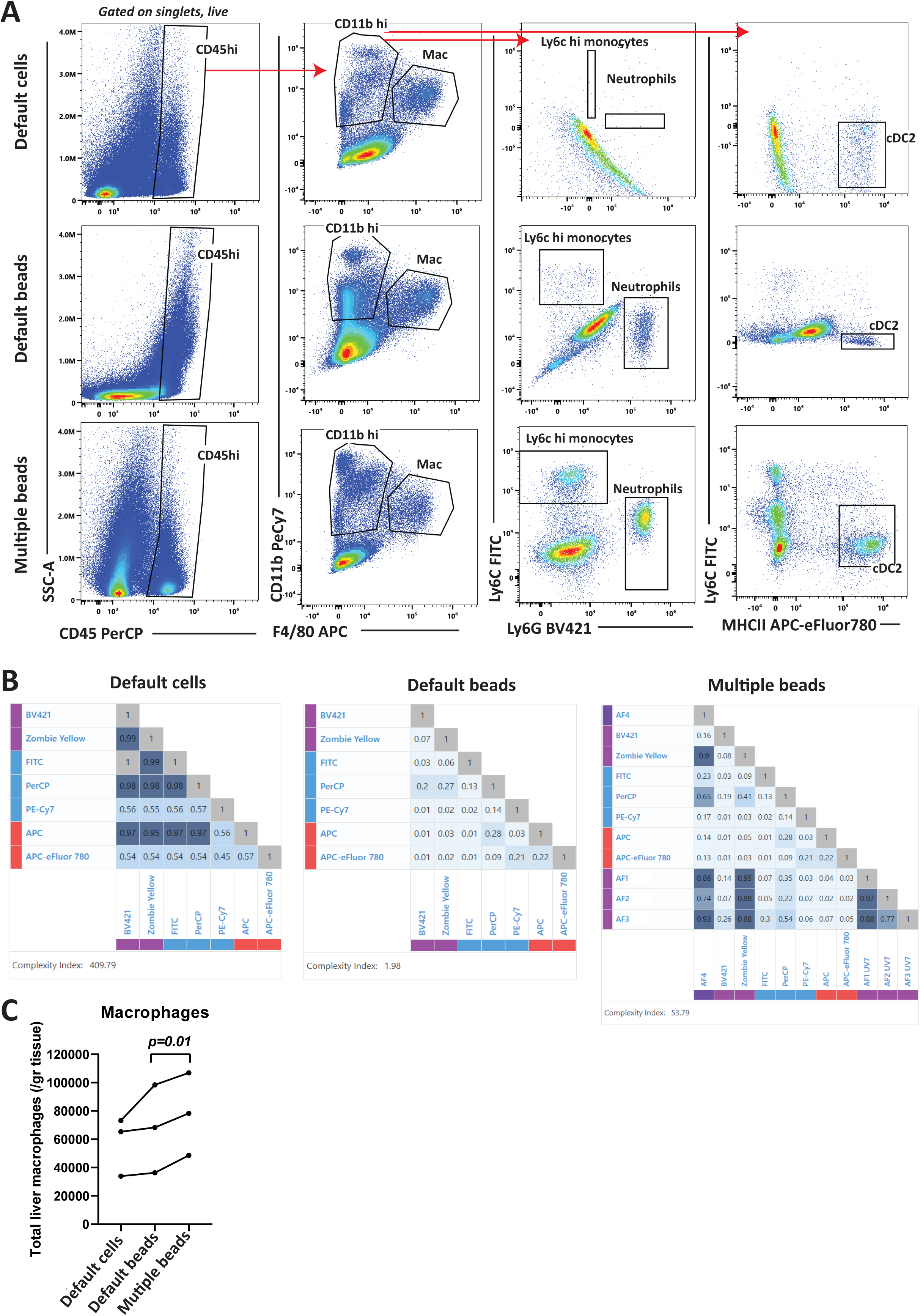
Default AF extraction in spectral flow cytometry in liver cells is not sufficient. A 7-marker panel was tested on the cell extraction of a healthy mouse liver. (A) shows gating strategies starting of CD45 cells (pre-gated on live, single cells), which were then separated into macrophages and other myeloid cells based on CD11b and F4/80. Myeloid cells (CD11b^hi^) were further divided in neutrophils, Ly6c^hi^ monocytes and classical dendritic cell (cDC) subsets by Ly6C, Ly6G, and MHCII expression. Unmixing was either performed by using liver cells as reference controls and using default AF extraction, by using bead reference controls, or by using bead reference controls in combination with multiple AF extraction. (B) Similarity matrix corresponding to the different approaches displayed in panel A. (C) Quantification of absolute number of macrophages per gram of liver tissue according to the different approaches displayed in panel A in the same samples (*n*=3). Mac = macrophages

To resolve these unmixing errors, we took a two-step approach. First, we switched to the use of beads for all fluorochromes except the live/dead stain. While using isolated liver cells as reference controls, the similarity index (SI) showed almost complete similarity between the fluorochromes FITC and BV421, which theoretically display different emission spectra (Figure 1B). This result highlights the inability to identify correct fluorochrome signatures within samples with high and heterogenous AF signals. The use of flow cytometry beads for reference controls instead of cells corrected this issue (Figure 1B).

Secondly, as the AF spectrum of the unstained liver cell suspension revealed the presence of at least two different AF spectra (Supplemental Figure 1A, arrows), we performed multiple AF identification manually. Hereto, we identified and extracted the single AF spectra by following a previously described protocol (13). Briefly, using an NxN strategy, we created bi-dimensional plots for each combination of detectors. We then selected raw channel combinations across the entire spectrum of detectors that best separated the populations, allowing us to gate different AF spectra. Finally, unique AF spectra were identified based on the SI. An SI threshold of 0.98 was used, above which two spectral signatures were considered insufficiently distinct to be resolved, as recommended by the manufacturer. It should be noted that after finishing the analysis of the data presented in this paper the Autofluorescence Explorer module in SpectroFlo® became available which now applies the same method. We observed a total of 4 unique spectra in the healthy livers (AF Control (Con)-1-4) (Supplemental Figure 1B-C and Figure 1B, right panel). We subsequently extracted the cells with the brightest signal for each of the identified single AF spectra (Supplemental Figure 1B) and added to the unmixing as unique fluorescent signatures. Only when using beads combined with the multiple AF extraction, we observed clearly defined populations of neutrophils and monocytes (Figure 1A). Importantly, the number of macrophages (CD11b^int^ F4/80^hi^) significantly increased with this method compared to using beads and default AF extraction, even though no direct unmixing issues were initially observed for the APC and Pe-Cy7 labels for F4/80 and CD11b (Figure 1C). Supplemental Figure 2 shows the bivariate plots of channels exhibiting high AF (Supplemental Figure 2A) and of the individual fluorochromes with overall AF in the whole liver sample used (Supplemental Figure 2B) before and after multiple AF extraction and using bead reference controls, showing improved signals. Some small unmixing errors remained, but these can be corrected manually by spillover correction in SpectroFlo®. Data shown are without manual corrections.

### A steatotic liver displays different autofluorescence than a healthy liver

Following this proof of principle, we moved to a 16-marker panel (Table 1, Figure 2) to identify all known macrophage populations in healthy liver and MASLD as well as other myeloid cells, i.e. neutrophils, eosinophils and several dendritic cell subsets. We successfully applied this on a healthy liver sample (Figure 2). We next investigated whether AF was similar in a steatotic liver by studying the liver of *db/db* mice. Due to leptin-deficiency, these mice inherently develop liver steatosis even on a standard chow diet (14). Interestingly, the average spectral signature of unstained cells isolated from the steatotic liver was distinct from that of a healthy liver (Supplemental Figure 3A). We then applied the same single AF spectra identification protocol for the steatotic liver as for the healthy liver. With this approach, we distinguished four populations with different AF properties compared to control livers (Supplemental Figure 3B-C) (AF Steatotic liver (SL)-1-4). AF SL-2-4 displayed high similarity to control AF spectra (SI>0.98). AF SL-1 was distinct from any of the AF signatures identified in the healthy liver (Supplemental Figure 3D).

**Figure 2.**
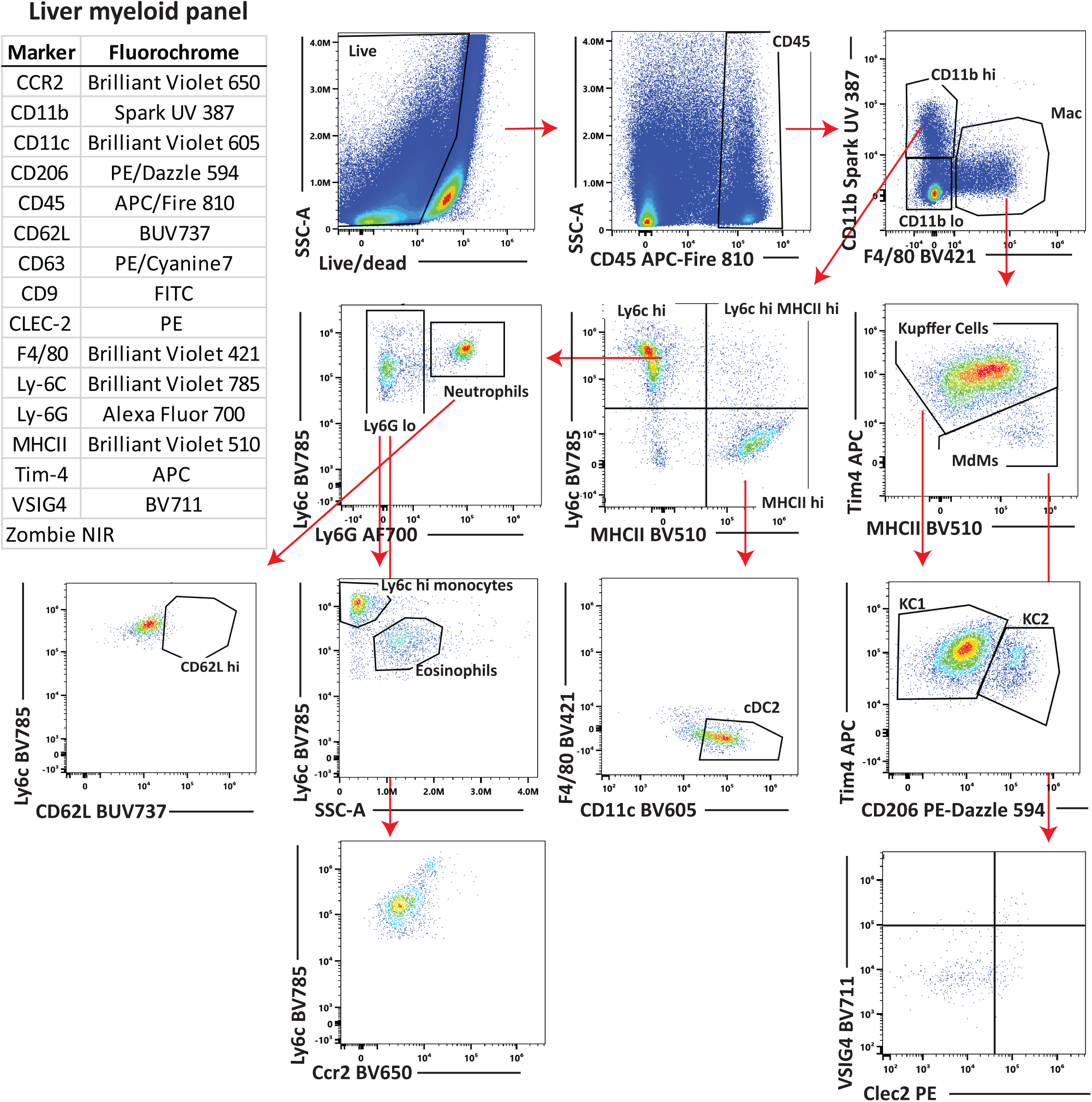
16-marker panel for liver myeloid cells in the liver. Overview of the 16-marker panel and gating strategy of the main cell populations using a 16-marker panel for liver myeloid cells in a healthy liver. Mac = macrophages. MdMs = Monocyte-derived Macrophages, cDC2 = type 2 conventional Dendritic Cells

**Table 1.** 16-marker panel for mouse liver macrophages.

| Target | Conjugate | Clone | Supplier | Catalog Number | Dilution |
| --- | --- | --- | --- | --- | --- |
| CD11b | Spark UV 387 | M1/70 | Biolegend | 101292 | 1:200 |
| Ly-6G | Alexa Fluor 700 | 1A8 | Biolegend | 127622 | 1:200 |
| Tim-4 | APC | RMT4-54 | Biolegend | 130022 | 1:200 |
| CD45 | APC/Fire 810 | 30-F11 | Biolegend | 103174 | 1:200 |
| F4/80 | Brilliant Violet 421 | BM8 | Biolegend | 123132 | 1:50 |
| MHCII | Brilliant Violet 510 | M5/114.15.2 | Biolegend | 107636 | 1:200 |
| CD11c | Brilliant Violet 605 | N418 | Biolegend | 117334 | 1:200 |
| CCR2 | Brilliant Violet 650 | SA203G11 | Biolegend | 150613 | 1:100 |
| Ly-6C | Brilliant Violet 785 | HK1.4 | Biolegend | 128041 | 1:200 |
| CD9 | FITC | MZ3 | Biolegend | 124808 | 1:100 |
| CLEC-2 | PE | 17D9 | Biolegend | 146104 | 1:200 |
| CD63 | PE/Cyanine7 | NVG-2 | Biolegend | 143910 | 1:100 |
| CD206 | PE/Dazzle 594 | C068C2 | Biolegend | 141732 | 1:50 |
| CD62L | BUV737 | MEL-14 | BD | 612833 | 1:100 |
| VSIG4 | BV711 | 17C9 | BD | 749502 | 1:200 |
| Zombie | NIR | - | Biolegend | 423105 | 1:250 |

### Incomplete AF extraction results in the inability to differentiate macrophage subsets

We next tested the impact of using different combinations of AF signatures on unmixing by removing the unique AF spectra one by one from the unmixing. Interestingly, removing the shared AF spectrum Con-4/SL-4 resulted in the inability to discriminate the two separate populations of KCs, i.e. CD206^low^ KCs and CD206^hi^ KCs (Figure 3A), both in the healthy liver and the steatotic liver. These two populations of KCs have previously been described and are termed KC1 (CD206^lo^) and KC2 (CD206^hi^) (15). KC1 and KC2 are also characterized by different expressions of CD63 and CD9, which are included in our panel. Indeed, the CD206^hi^ KCs were also characterized by a higher expression of both CD63 and CD9 (Figure 3B). For CD63, the same issue was observed as for CD206, i.e. two separate cell populations were only observed with AF Con-4/AF SL-4 added to the unmixing.

**Figure 3.**
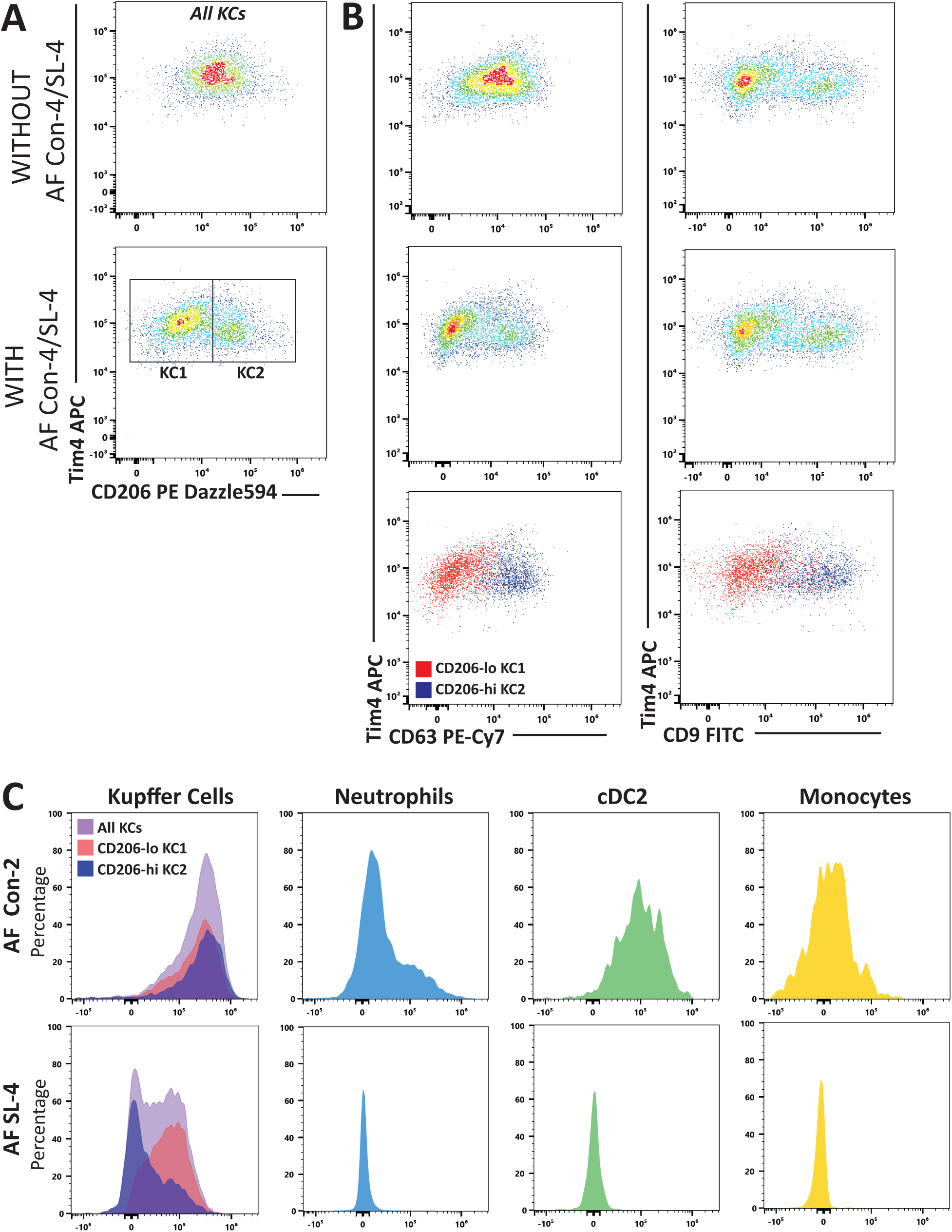
AF spectrum is essential for distinguishing Kupffer Cell subpopulations. (A) Addition of the AF signature AF Con-4/SL-4 to the unmixing was essential for the separation of CD206-lo KC1 and CD206-hi KC2 with the total KC population. Gating for KCs done as shown in Figure 2. (B) KC1 and KC2 also displayed differential expression of CD63 and CD9. (C) Histograms of the signal of AF signatures in different KC populations as well as neutrophils, cDC2 and monocytes (gating of populations as shown in Figure 2), revealing AF Con-4/SL-4 to be specific for KC1. KC = Kupffer Cell, cDC2 = type 2 conventional Dendritic Cells

With the multiple AF extraction, the unique AF spectra identified are designated as fluorescence tags in the experiment. This allows us to treat these AF spectra like other fluorochromes in the experiment. Thus, we were able to analyze the emission of these AF signatures by the different cell populations. Liver macrophages expressed all identified AF spectra, further confirming the relevance of all these spectra for proper unmixing. AF Con-4/AF SL-4 was essential for separating KC1 and KC2 cells, suggesting that emission of this AF signature differs between these two subsets of KCs. Indeed, AF Con-4/AF SL-4 was specific for CD206^lo^ KC1 as opposed to CD206^hi^ KC2 or other immune cells (Figure 3C, bottom). In contrast, the AF spectrum Con-2 for example was emitted by all KCs and was highly specific for KCs in general (Figure 3C, top). Thus, incomplete AF extraction resulted in the inability to distinguish between these smaller macrophage subpopulations.

### Separate AF handling biases cell quantification between healthy and MASH livers

The steatotic livers of Db/Db mice did not exhibit monocyte-derived macrophage (MdM) infiltration. Therefore, we next tested the 16-marker panel and AF correction strategy outlined above on livers from male mice fed a MASH diet or a matched control diet (healthy) for 10 weeks. We also performed multiple AF extraction in both the healthy and MASH diet livers and found 5 spectra in each condition. Comparison of these 10 AF spectra revealed 7 unique AF spectra (SI<0.98) across the two conditions; 3 shared AF spectra (AF Healthy (H)-1-3/AF MASH (M)-1-3), 2 unique spectra in the healthy liver (AF H-4,-5) and 2 unique spectra in the MASH liver (AF M-4,-5) (Supplementary Figure 5A).

A practical question now arose on how to accommodate these differences in AF between experimental conditions during unmixing. Two approaches are possible: (i) separate unmixing per condition using their own AF sets, or (ii) combined unmixing that includes all AF components deemed essential across conditions. We hypothesized that separate unmixing risks non-comparable quantification. To test this, we performed spectral unmixing for each group using either its own AF spectra or the other group’s AF spectra (healthy unmixed with healthy vs MASH AF, and vice versa) (Figure 4B, Supplementary Figure 4). Quantification of total immune cells (CD45^+^), total macrophages and ly6c^hi^ monocytes was significantly different comparing unmixing with the different set of AF spectra, predominantly in the control group (Figure 4A) without obvious visual unmixing artifacts for either unmixing strategy (Supplementary figure 4). This demonstrates that using different AF matrices between groups can bias cell frequency quantifications.

**Figure 4.**
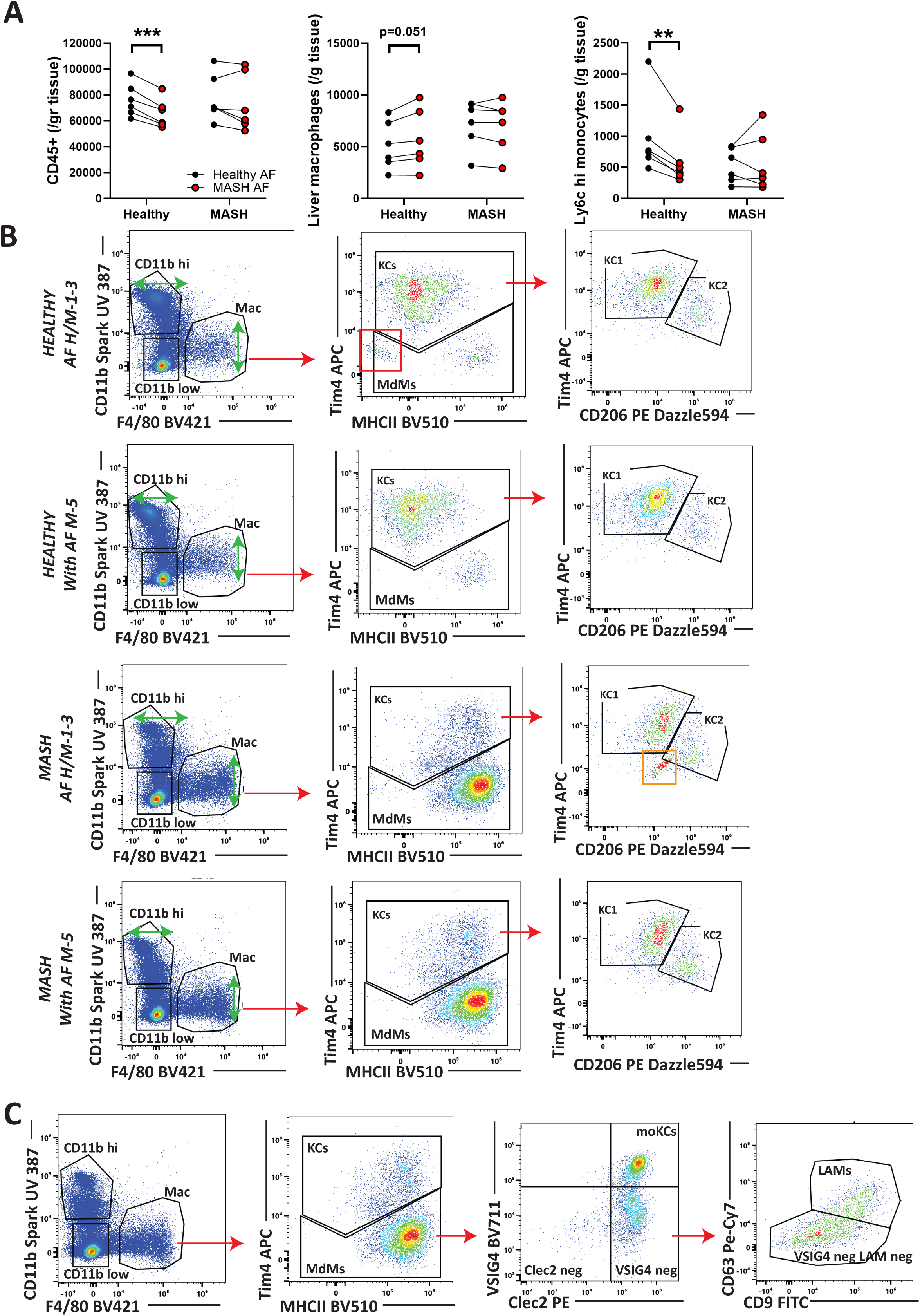
Separate AF handling biases cell quantification between healthy and MASH livers. (A) Mice were fed a 10-week MASH or control diet (healthy) (*n*=6/group). Unmixing was done using healthy AF versus MASH AF spectra in both conditions. CD45^+^, total macrophage and Ly6c^hi^ cells were quantified. Gating is similar to Figure 2. Paired t-test was used, ** p<0.01, *** p<0.001. (B) AF signature AF M-5 improves for unmixing of liver macrophages in healthy and MASH livers in addition to shared AF spectra H/M-1, −2, −3. Red and orange boxes indicate unmixing inaccuracies and green arrows indicate spread of populations. (C) shows the macrophage populations in a liver from a mouse on a 10-week MASH diet, including monocyte-Kupffer Cells (moKCs) and lipid-associated macrophages (LAMs). Unmixing was performed using the AF spectra from the MASH liver and gating was similar to Figure 2. Mac = Macrophage, KC = Kupffer Cell, MdM = Monocyte-derived monocytes, moKCs = monocyte Kupffer Cells, LAMs = Lipid-Associated Macrophages, M = MASH, H = Healthy.

Therefore, we next tested a combined unmixing strategy to ensure robustness of results between experimental groups. Because adding too many AF components increases data spread and reduces resolution, we first identified which AF spectra were essential for accurate macrophage unmixing in both the healthy and MASH liver. We started by performing unmixing with all unique AF spectra, which resulted in suboptimal unmixing in predominantly the healthy liver seen by an aberrant Tim4^−^/MHCII^−^ population in the F4/80^hi^ cells (Supplementary Figure 5B, red boxes). We then assessed the relevance of the three shared AF spectra by removing these AF spectra one by one from the unmixing in both the healthy and MASH livers. This resulted in worsening of the aberrant Tim4^−^/MHCII^−^ population in the F4/80^hi^ cells in the healthy liver and appearance of such population in the MASH liver (Supplementary Figure 5B, red boxes) as well as increased spreading (Supplementary Figure 5B, green arrows) indicating that these shared spectra are indispensable.

Inclusion of only these three shared AF spectra did not result in optimal unmixing, as seen by abnormal cell populations in the healthy and MASH liver (Figure 4B, red and orange boxes). Next, we evaluated whether adding condition-unique AF spectra further improved unmixing by adding them one-by-one to the unmixing. Notably, inclusion of AF M-5 not only eliminated the abnormal cell populations in the MASH but also healthy liver (Figure 4B, red and orange boxes) and reduced spread within CD11b^hi^ and macrophage populations in both conditions (Figure 4B, green arrows). Inclusion of other condition-specific AF spectra did not notably change unmixing (data not shown).

As expected, MASH development led to the appearance of a large number of MdMs (Figure 4C, gating similar to healthy liver in Figure 2) that are not found in a healthy liver (Figure 2) or the steatosis-only liver (data not shown). Importantly, we could positively identify LAMs using CD9 and CD63, moving beyond prior reliance on negative gating in conventional flow cytometry (11). Identification of LAMs in the liver by flow cytometry has remained challenging, and this data indicates that detailed correction for AF may be a key prerequisite for accurate LAM identification.

Together, these results support a combined unmixing strategy built from (i) the shared AF spectra required for accurate macrophage resolution and (ii) select condition-unique spectra (e.g. AF M-5) that demonstrably improve unmixing and resolution without inflating spread. Using a common AF basis across groups can preserve quantitative comparability. Our data also suggests that manual AF extraction may have missed a relevant AF component in healthy liver. This may also explain why in our test of Figure 4A predominantly cell quantification of the healthy liver was affected by using MASH AF spectra. Cross-condition AF pooling may better capture the full AF landscape in complex tissues like liver.

## Discussion

We aimed to develop a spectral flow cytometry approach to identify and analyze all macrophage subpopulations in healthy and MASH murine livers. As the liver displays high and heterogeneous AF, i.e. the presence of multiple AF signatures, multiple AF extraction was essential. Although default AF extraction during unmixing was able to identify a defined macrophage population, default AF extraction resulted in a significant underestimation of the number of macrophages compared to analysis with multiple AF extraction. Importantly, this was not associated with any signs of incorrect unmixing, highlighting the danger of unknown measurement errors with improper AF extraction for the correct quantification of tissue cells using spectral flow cytometry.

It has previously been described that KCs in the healthy liver can be divided into two distinct populations, termed KC1 and KC2, which can be separated by expression of several markers including CD206 (15). However some doubts have been raised whether these populations actually exist (16). Here we demonstrated that CD206^lo^ KC1 and CD206^hi^ KC2 can emit distinct AF spectra and that adequate AF extraction was essential to distinguish KC2 from KC1 in our study. Importantly, this may hamper the separation of KC1 and KC2 in conventional flow cytometry experiments or spectral flow cytometry experiments without proper AF handling and may contribute to inconsistencies between studies on the identification of these two separate populations. A previous report showed that extracting AF spectra from macrophages in the lung can improve resolution. Rather than performing visualization of unstained events across N x N plots, they described an interesting approach using high dimensionality dimensions reductions to identify AF spectra within a given tissue. However, this approach failed to distinguish subtle changes within the macrophage subset (2). Since our data indicate that two KC subsets can display slightly different AF spectra, it seems advisable to apply different approaches for the identification of such resembling spectra. Our results underscore not only that accounting for subtle, cell type-specific AF is essential for accurate resolution of closely related immune subsets, but it will also be essential for ensuring reproducibility and comparability across studies. This is also demonstrated by the fact that with our approach we are now able to identify LAMs based on known signature markers where previously LAM identification by flow cytometry relied on negative gating (11).

A key methodological challenge we encountered was the presence of partially distinct AF spectra between experimental conditions, e.g. healthy and MASH livers. Protocols for multiple AF extraction in case of heterogeneous AF have been described previously (13), but they typically do not account for changes in AF signatures between experimental conditions. While a straightforward approach is to unmix each experimental group with its own AF spectra, our data show that this approach leads to substantial differences in the quantification of major immune cell populations, even in the absence of overt unmixing artifacts. This can lead to erroneous conclusions about differences between experimental groups. A more robust approach to be considered is to include condition-unique AF spectra in the unmixing of all groups, thereby ensuring comparability.

Importantly, considering all AF spectra of all conditions in the unmixing can induce overcorrection and loss of data resolution. As advised by the manufacturer, including more than five AF spectra in the unmixing will inevitably lead to increased spread and loss of data resolution. Therefore, it is essential to identify those impactful AF spectra that are essential for identification of the immune cells of interest in every experimental condition. A detailed method for identifying relevant AF spectra has been described by Kharaz et al (5). When using a method of combined unmixing of all conditions, pooling of unstained samples of experimental conditions can also be performed reducing the need to analyze and compare AF in all unstained samples. However, insights in relative AF expression between experimental conditions is lost in this case. Importantly, we encountered a situation where addition of a MASH AF spectrum also improved the unmixing of the healthy liver. This finding suggests that manual AF extraction may fail to capture all relevant AF components in complex tissues and that inclusion of spectra from multiple conditions can compensate for such limitations. This also highlights the need for automated methods to reliably identify and extract distinct AF profiles in complex samples such as liver-derived cell suspensions. In our MASH study we have chosen for combined unmixing, i.e. include all essential AF spectra from control and MASH liver in the unmixing of all samples because of 1) the high complexity of these samples and high chance of missing essential spectra in either experimental group and 2) our need to reliably compare absolute cell numbers as well as relative expression of activation markers on macrophages populations between experimental conditions to answer our research questions.

In conclusion, we show that appropriate AF extraction is essential for reliable identification and quantification of liver macrophages by spectral flow cytometry, particularly in complex tissues such as the MASH liver. When properly handled, full spectral flow cytometry enables robust high-dimensional immune profiling, which is critical for understanding macrophage behavior across health and disease and for advancing therapeutic strategies targeting hepatic macrophage subsets.

### Data Limitations and Perspectives

Our study has several limitations. First, AF extraction and selection was performed manually, which is subjective to operator bias and may not fully capture the complexity of AF profiles in liver suspensions. Automated and standardized AF identification methods will be required to reduce operator bias and improve reproducibility across laboratories. Second, although we chose a combined unmixing strategy to ensure comparability between healthy and MASH samples, we cannot exclude that this approach incorporates AF spectra not present in all samples. Objective criteria for defining the minimal AF set needed per experiment remain to be established. Third, while our data indicate that AF may contribute to inconsistencies in the reported separation of KC subsets, our study was not designed to resolve the biological debate on their existence, which will require validation strategies.

Looking ahead, condition-dependent AF variability is likely also relevant in other experimental systems, even simple settings as cell culture experiments, and may be influenced by technical factors such as enzyme cocktails or dissociation protocols. The field would benefit from systematic benchmarking of AF handling strategies, the development of automated AF extraction workflows, and guidelines for panel design and experimental set-up that integrate AF considerations from the onset. These steps will be essential to ensure robust, comparable high-dimensional flow cytometry data, particularly as more subtle immune cell subsets continue to be identified.

## Materials and Methods

### Animal model

8- to 12-week old male C57BL/6 mice and 10-week old male db/db mice (BKS.Cg-Dock7^m^+/+Lepr^db^J Charles River) were used. In addition, 10-week old male C57BL/6 mice were fed the Gubra-Amylin NASH diet (GAN) diet (E15746, SSniff diets) or matched control diet for 10 weeks. All performed experiments were approved by the Animal Experiments Committee of Maastricht University.

### Cell isolation and staining

After mice were euthanized, the liver was perfused manually with PBS via the portal vein. The liver was minced and transferred to a tube containing 15mL of RPMI (1640 GlutaMAX, Gibco) with Collagenase A (0.75 mg/ml; Sigma) and DNase (50 µg/ml; Sigma) and then incubated for 30 min at 37C° with frequent mixing. The liver digest was then passed through a 70-micron filter and washed with 15 mL of RPMI (GlutaMAX) supplemented with 10% FBS and 1% P/S. The cell mixture was centrifuged at 50xg for 3 min at 4°C. The supernatant was then transferred to a new falcon tube followed by centrifuging at 340xg for 7 min at 4°C. The supernatant was discarded. To lyse red blood cells, the pellet was resuspended in lysis buffer (0.1 mM EDTA, 0.84% NH_4_CL) and incubated for 5 min at room temperature. The cells were washed with 10mL of PBS and pelleted at 340xg for 7 min at 4°C. The supernatant was discarded, and cells were resuspended in 1mL of FACS buffer (1X PBS with 10 mM Na azide and 0.5% BSA in g/mL). To prepare the samples for flow cytometry, the cells were transferred to 1.5mL Eppendorf tube and pelleted at 650xg for 3.5 min at 4°C. For live/dead staining, the cell pellet was resuspended in 100µL of 1:250 Zombie Yellow or NIR and incubated for 15min, on ice, in the dark. The cells were then washed with 500µL of FACS buffer and re-pelleted at 650xg for 3.5 min at 4°C. To prevent non-specific antibody staining, pellets were incubated with 10µL of 1:10 Fc-block (in FACS buffer) on ice for 5 min. Following this the cells were incubated with 90µL of fluorochrome-conjugated antibodies (Table 1, Supplementary Table 1) for 45-60 min on ice and in the dark. Antibody dilutions were determined based on titration. Live-dead staining dilutions were based on titration including mixtures of live and dead cells. The stained cells were washed with 500µL of FACS buffer and re-pellet at the same speed at 4°C. The cells were resuspended in 300µL of FACS buffer and filtered for flow cytometry analysis.

### Spectral flow cytometry and analysis

All samples were measured on a Cytek Aurora full spectral cytometer equipped with 4 lasers, i.e. UV 355nm, violet 405 nm, blue 488 nm, and red 638 nm, and 54-channel detection (16UV, 16V, 14B, 8R). In all cases, manufacturer quality control programs were run and passed before acquisition of data. Acquisition, unmixing and analysis were performed using Cytek SpectroFlo® v3.0.1 software. AF extraction was done either via default extraction available in the SpectroFlo software or was done manually as described in (13). Fluorescence minus one (FMO) controls were performed for each antibody in order to set gate borders and assess AF.

## Abbreviations

MASH: Metabolic Dysfunction-Associated Steatohepatitis
AF: autofluorescence
MASLD: Metabolic dysfunction-Associated Steatotic Liver Disease
moKCs: Monocyte-Kupffer Cells
LAMs: Lipid-Associated Macrophages
KC: Kupffer Cell
MdMs: Monocyte-derived Macrophages
cDC2: type 2 conventional Dendritic Cells

## Funding information

SD is supported by a MSCA Marie Curie Postdoctoral Fellowship (MacTalk, 101063130). KW and SD are supported by a TKI-grant (translATe-NASH, LSHM202312)

## Conflict of interest statement

The authors declare no conflicts of interest.

## Data availability statement

The data that support the findings of this study are available from the author upon reasonable request.

## Ethics Statement of animal experiments

All mouse experiments were approved by the Animal Experiments Committee of Maastricht University (AVD10700202317035).

## Author contributions

**Sabine Daemen:** Conceptualization, Investigation, Funding acquisition, Writing original draft, Methodology, Review & editing, Visualization

**Panos Barlampas:** Investigation, Review & editing

**Xiaodi Zhang:** Investigation, Review & editing

**Casper Schalkwijk:** Supervision, Review & editing, Resources

**Kristiaan Wouters:** Conceptualization, Review & editing, Supervision, Resources, Methodology, Funding acquisition

## Notes

### Competing Interest Statement

The authors have declared no competing interest.

