## Supplemental Material for "Managing Autofluorescence in Spectral Flow Cytometry for Macrophage Identification in the Liver"

**Supplementary information Daemen et al.**

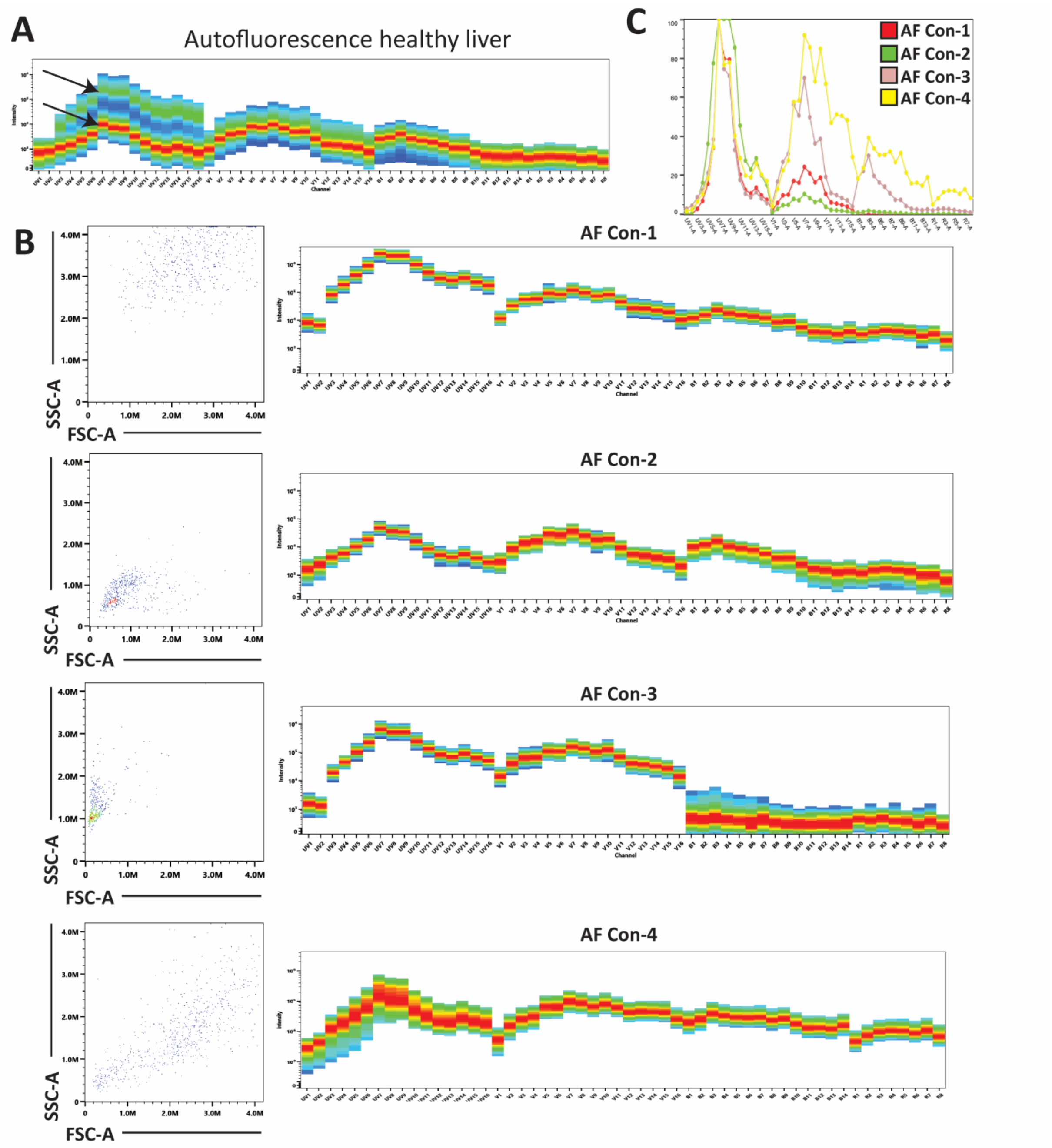


**Supplemental Figure 1 Autofluorescence spectra in the healthy murine liver.** (A) shows the raw AF signal of unstained healthy liver cells in all detectors. Arrows indicate the presence of at least two different AF spectra. (B) shows FSC and SSC plots and the AF signatures of the four populations identified in the healthy mouse liver. (C) shows an overlay of the identified AF spectra. AF = autofluorescence, Con = control

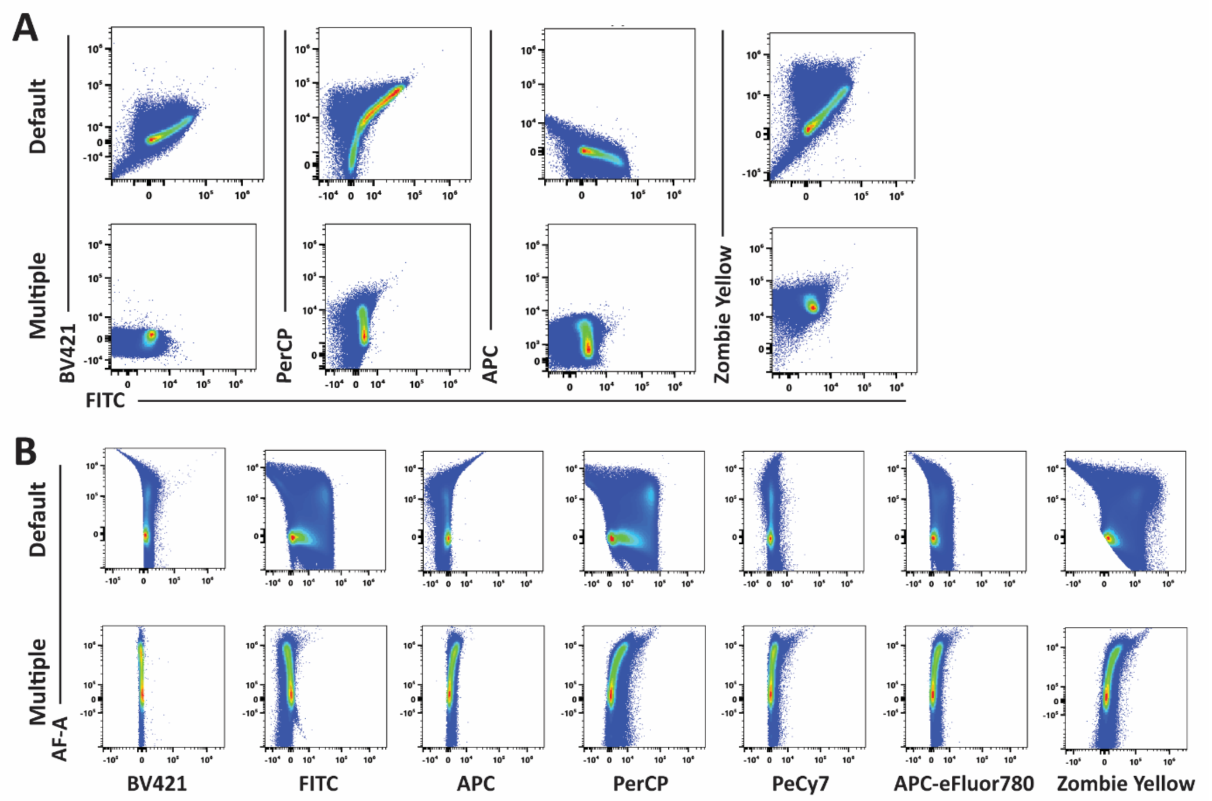


**Supplementary Figure 2 Multiple AF extraction corrects unmixing errors.** (A) Bivariate plots of channels exhibiting high autofluorescence before and after multiple AF extraction and using bead reference controls. (B) Bivariate plots of individual fluorochromes with overall AF in the whole liver sample before and after multiple AF extraction.

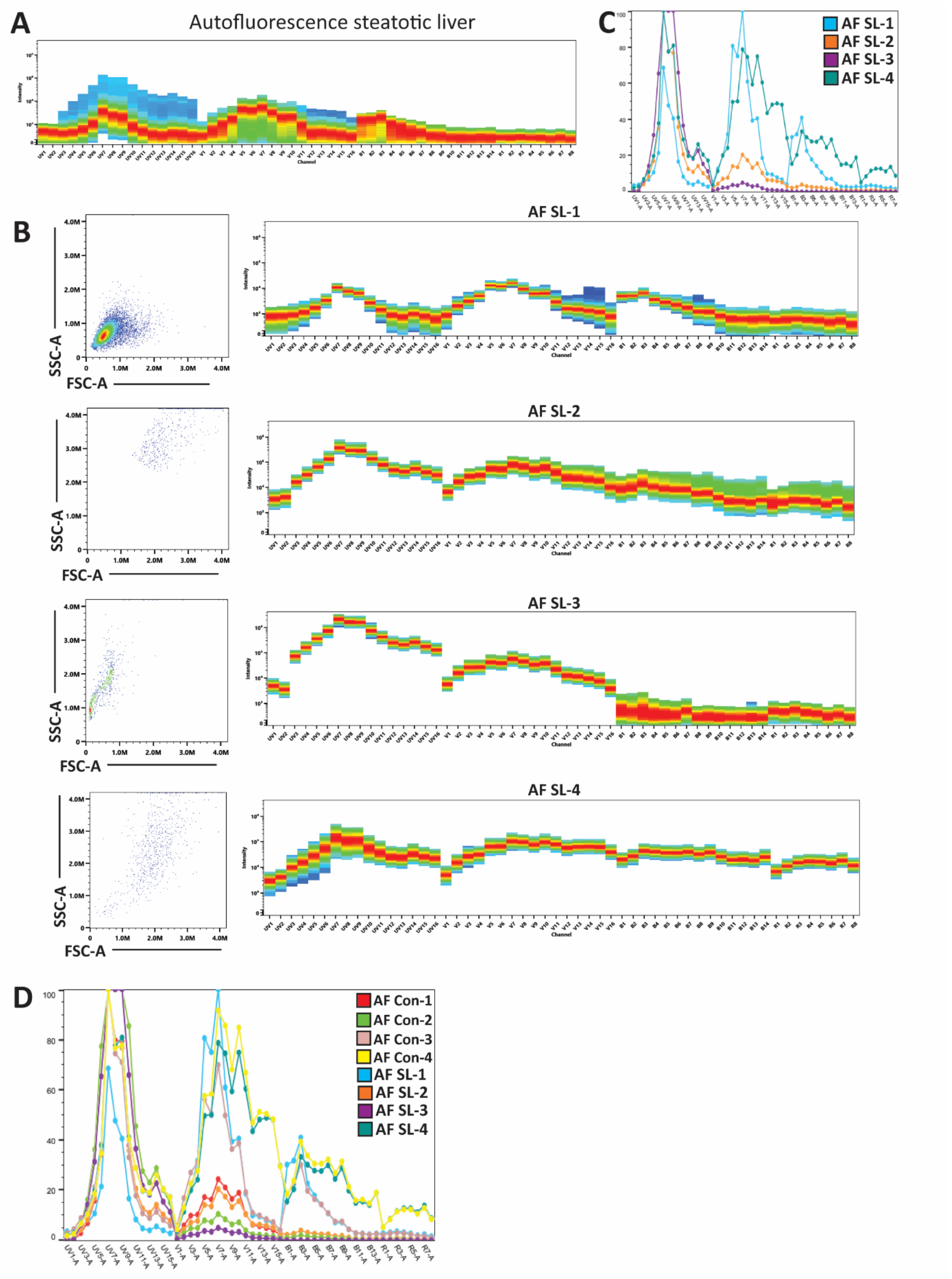


**Supplementary Figure 3 AF differs between a healthy liver and a steatotic liver.** (A) AF of a liver cells extracted from a steatotic liver of a *db/db* mouse. (B) displays FSC-SSC scatter plots and AF signatures of the populations with different AF in livers of Db/Db mice. (C) shows an overlay of the identified AF spectra. (D) shows an overlay of all AF signatures identified in both healthy and steatotic livers. AF = autofluorescence, SL = steatotic liver

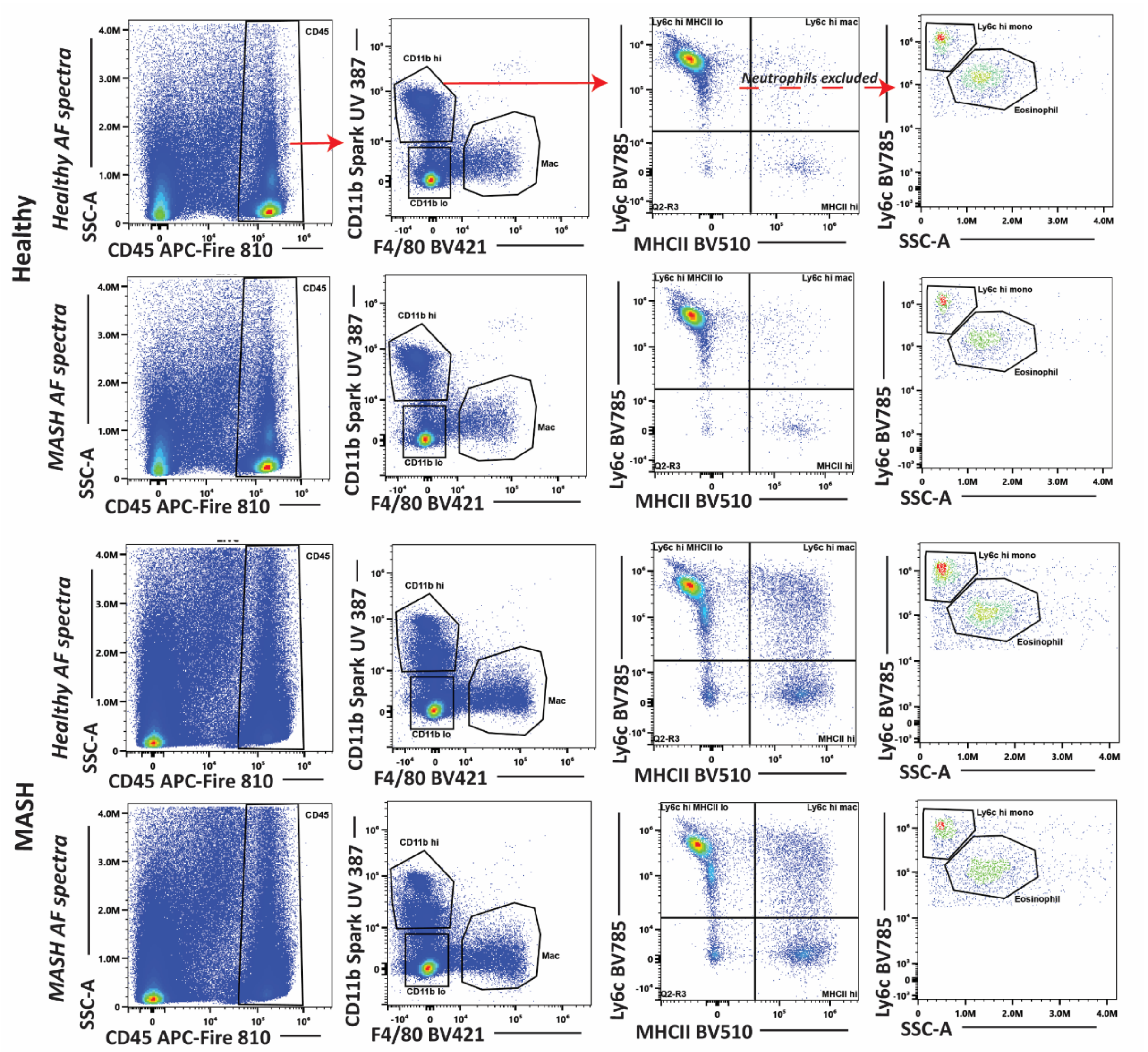


**Supplementary Figure 4 Unmixing with healthy or MASH AF spectra does not visually change unmixing.** Healthy and MASH livers were unmixed using either all healthy AF spectra or MASH AF spectra.

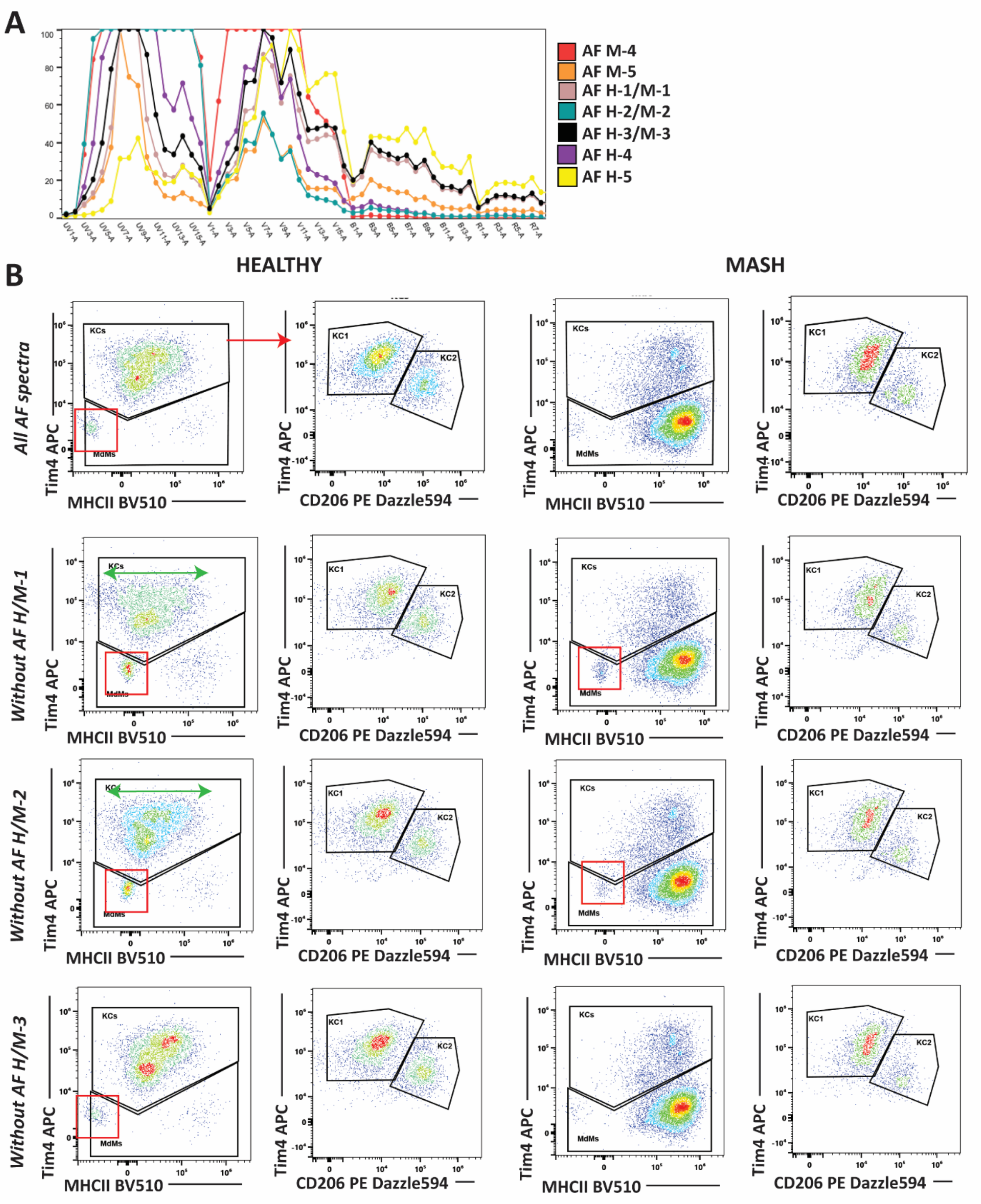


**Supplementary Figure 5 Three shared AF signatures (AF H/M 1-3) between healthy and MASH livers are essential for unmixing.** (A) All unique AF spectra identified in healthy (H) and MASH (M) liver, including three shared AF spectra and two unique AF spectra in each condition. (B) Removal of the shared AF signatures from unmixing induces macrophage unmixing inaccuracies which are highlighted by the red boxes and increased spreading by green arrows. KC = Kupffer Cell, MdM = Monocyte-derived monocytes

**Supplementary Table 1. 7-marker test panel for mouse liver myeloid cells**

| Target | Conjugate | Clone | Supplier | Catalog Number | Dilution |
|---|---|---|---|---|---|
| Ly6G | BV421 | 1A8 | Biolegend | 127628 | 1:100 |
| Ly6c | FITC | 1G7.G10 | MACS | 130-123-285 | 1:100 |
| F4/80 | APC | BM8 | Biolegend | 123116 | 1:100 |
| CD45 | PerCP | 30-F11 | Biolegend | 103130 | 1:100 |
| CD11b | PeCy7 | M1/70 | BD | 561098 | 1:100 |
| MHCII | APC-eFluor780 | M5/114.15.2 | Invitrogen | 47-5321-80 | 1:100 |
| Zombie | Yellow | | Biolegend | 423103 | 1:100 |
